# Immunofocusing humoral immunity potentiates the functional efficacy of the AnAPN1 malaria transmission-blocking vaccine antigen

**DOI:** 10.1101/2020.11.29.402669

**Authors:** Nicole G. Bender, Prachi Khare, Juan Martinez, Rebecca E. Tweedell, Vincent O. Nyasembe, Borja López-Gutiérrez, Abhai Tripathi, Dustin Miller, Timothy Hamerly, Eric M. Vela, Randall F. Howard, Sandrine Nsango, Ronald R. Cobb, Matthias Harbers, Rhoel R. Dinglasan

## Abstract

Malaria transmission-blocking vaccines (TBVs) are a critical tool for disease elimination. TBVs prevent completion of the developmental lifecycle of malarial parasites within the mosquito vector, effectively blocking subsequent infections. The mosquito midgut protein Anopheline alanyl aminopeptidase N (AnAPN1) is the leading, mosquito-based TBV antigen and structure-function studies have identified two Class II epitopes that induce potent transmission-blocking (T-B) antibodies. Here, we functionally screened new immunogens and down-selected to the UF6b construct that has two glycine-linked copies of the T-B epitopes. We established a process for manufacturing UF6b and evaluated in outbred female CD1 mice the immunogenicity of the preclinical product with the human-safe adjuvant Glucopyranosyl Lipid Adjuvant in a liposomal formulation with saponin QS21 (GLA-LSQ). UF6b:GLA-LSQ was immunogenic and immunofocused the humoral response to one of the key T-B epitopes resulting in potent T-B activity and establishing UF6b as a prime TBV candidate to aid in malaria elimination and eradication efforts.

## Introduction

Malaria is responsible for about 405,000 deaths each year, predominantly among children under the age of five in Sub-Saharan Africa^1^. To achieve malaria elimination and eradication, we must leverage concerted approaches to reduce clinical disease and prevent new cases. RTS,S (Mosquirix™), the most advanced malaria vaccine that has been rolled out in selected African countries, reduces disease burden for individuals, but alone it cannot ultimately support malaria eradication since it does not completely prevent malaria transmission^2–4^.

The *Plasmodium* parasites responsible for causing malaria have an obligatory developmental cycle in the female *Anopheles* mosquito vector, which begins when a mosquito takes up male and female *Plasmodium* gametocytes in a blood meal from infected humans. Within the mosquito bloodmeal bolus, the gametocytes transform to male and female gametes that combine to form motile zygotes (ookinetes), which must first attach to the midgut epithelium followed by enterocyte traversal to form an oocyst just beneath the midgut lamina. Within two weeks, mature oocysts rupture releasing thousands of sporozoites that are passively transported to the salivary glands. Malaria parasite transmission is completed once the infectious mosquito injects salivary fluid containing sporozoites while taking a blood meal from a naïve human host.

Transmission-blocking vaccines (TBVs) work by preventing parasite sporogonic development in the mosquito, thereby disrupting human-mosquito-human transmission (**Fig. 1a**)^2–5^. As a result, they also have the added benefit of preventing the spread of parasites that have developed drug resistance^2,6^. As a complementary intervention to RTS,S, TBVs may extend the utility of the RTS,S vaccine by reducing parasite breakthrough transmission in vaccinated populations^2–7^. While several TBVs that target *Plasmodium* proteins have already been tested in the clinic, parasite-centric vaccines are inherently species-specific, requiring the production and validation of several malaria vaccine modalities^2,3^. In contrast, a pan-malaria TBV, which targets a conserved mosquito protein that acts as a parasite receptor in the mosquito midgut, may address the limitations inherent in parasite-centric vaccines.

**Fig. 1.**
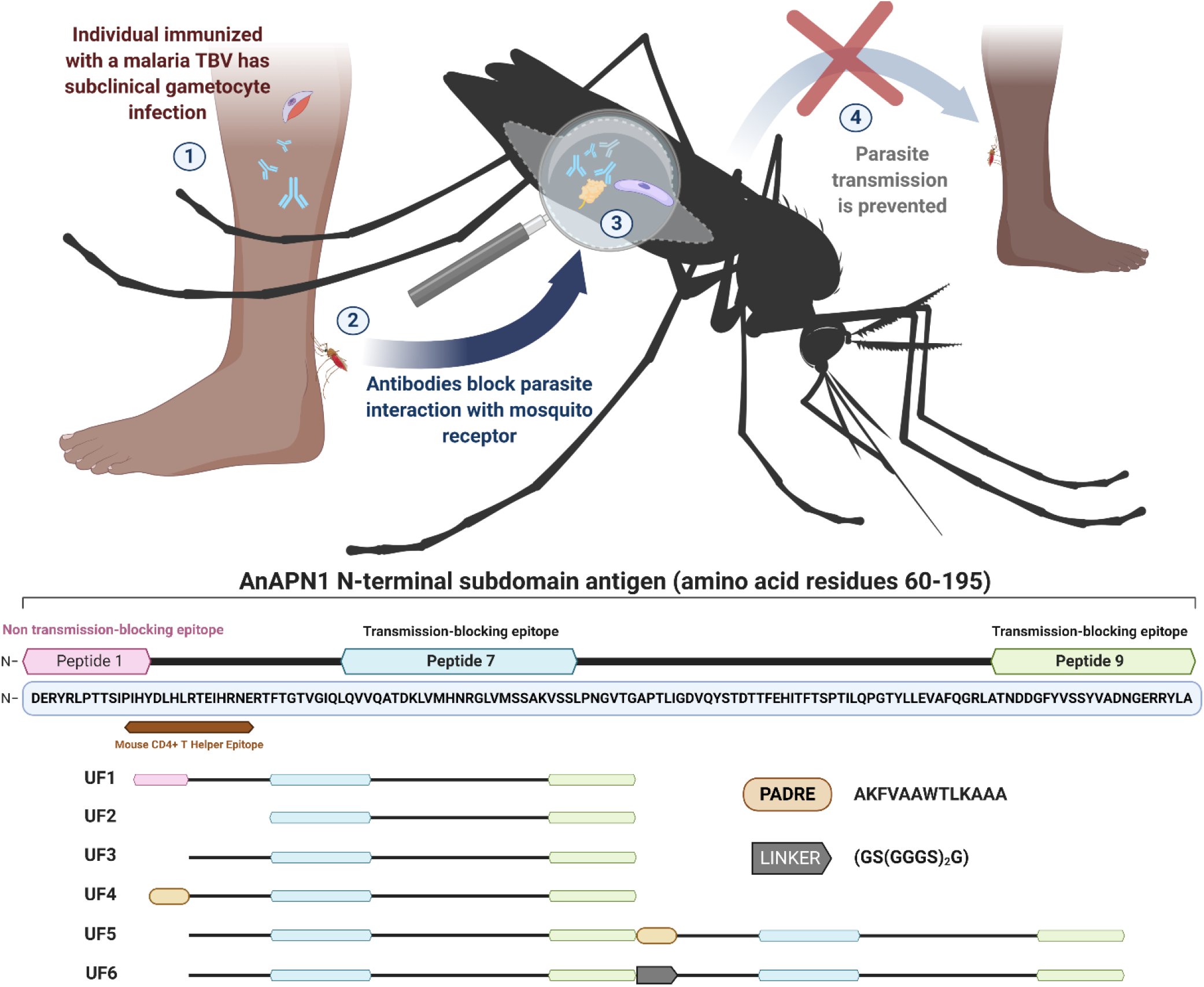
Design of AnAPN1 Transmission-Blocking Vaccine (TBV) constructs. **(Top)** Schematic depicting malaria mosquito-based TBV mode of action, wherein antibodies generated in a human host ① are taken up by the mosquito ③ and prevent parasite (purple)-protein receptor (yellow) interaction ② and subsequent development within the mosquito, effectively blocking the spread of parasites within a community ④. **(Bottom)** AnAPN1 subdomain antigen design. Several second-generation immunogen constructs of AnAPN1 were developed based on structural and functional information for peptides 1, 7, and 9 (note: peptide locations were not assigned left to right). Location of a Mouse T Helper Epitope (brown) identified through the Epitope Identification Suite (Merck Research Labs) is indicated7, which overlaps with the carboxy-terminal side of peptide 1. AnAPN1 constructs; UF1: Original AnAPN1 design, UF2: Mouse T Helper Epitope Deletion, UF3: Peptides 2-9 only, UF4: [PADRE: AKFVAAWTLKAAA] + Peptides 2-9 (UF3), UF5: UF3 + PADRE + UF3 construct, UF6: UF3+[Glycine Linker: GS(GGGS)_2_G] + UF3. (Protein domains are not drawn to scale). See **table S1** for additional details.

The mosquito midgut surface protein Anopheline Alanyl aminopeptidase N (AnAPN1) is an advanced, pan-malaria mosquito-based TBV candidate^7^. Antibodies raised against the N-terminal domain of this protein were shown to prevent the *Plasmodium falciparum* ookinete from traversing the midgut wall and developing into oocysts, blocking further development and transmission of the parasite^7–10^. Our previous studies demonstrated that anti-AnAPN1 antibodies can completely (100%) block transmission of naturally circulating *P. falciparum* in Cameroon^7–10^. By solving the crystal structure of AnAPN1, we mapped two conformational and protective transmission-blocking (T-B) epitopes, peptides 7 and 9, and a non-protective, “decoy” peptide 1 epitope^7,8^. Considering that a monoclonal antibody to Peptide 7 alone^8^, and polyclonal antibodies that target only Peptide 9 can both result in potent transmission-blocking activity^7^, a vaccine formulation that promotes a highly focused and functional humoral response in humans to either peptide epitope would be considered successful. Peptide 1 includes a mouse CD4+ T cell epitope, allowing it to act as an “immune decoy” and elicit a significant humoral response with a robust but non-protective antibody titer^8^. Therefore, removing this immune-decoy epitope from the original 125 amino acid subdomain design of the AnAPN1 TBV (125 amino acid recombinant AnAPN1)^7,8^ may prevent the induction of an unintended and a predominant humoral response to peptide 1, and consequently improve T-B activity. We hypothesized that altering the immunogen design would focus ideally the humoral immune response to both of the critical peptide epitopes (peptide 7 and 9) and potentiate T-B activity, essentially requiring lower antibody titers to be elicited in the mammalian host while still achieving potent T-B activity. Here, we report the results of an immunological and functional screen in mice of new AnAPN1 constructs that do not contain the mouse immunodominant peptide 1 epitope, and are capable of immunofocusing the vertebrate humoral response to at least one of the critical T-B epitopes, peptide 9. In this study, we not only demonstrated the marked immunogenicity and T-B functionality of the selected second-generation AnAPN1 construct (UF6), but also identified a potent formulation of the preclinical product and purification process for a tag-free immunogen (UF6b) with the human-safe Glucopyranosyl Lipid Adjuvant in a liposomal formulation with saponin QS21 (GLA-LSQ) adjuvant^9,11^. Overall, our results highlight UF6b:GLA-LSQ as a strong TBV candidate that may be a critical component in the fight to eliminate if not eradicate malaria.

## Results

### Down-selection to the UF6 immunogen construct

Previous studies on the T-B activity of anti-AnAPN1 IgG together with data on the protein structure revealed that there are multiple epitopes on the AnAPN1 protein, some of which contribute to T-B activity, while others, such as peptide 1 do not^7,8^. Hence, we used these data to inform the design of new immunogen variants to specifically present to the immune system only the relevant T-B domains of AnAPN1, thereby effectively avoiding undesired immunodominant epitopes. Using a wheat germ cell-free protein expression system, we prepared as a reference standard, the original AnAPN1 antigen construct (UF1), along with five new antigen candidates for the screening study (**Fig. 1b**, **fig S1a,b)**. A UF2 antigen was constructed based on UF1 without peptide 1, the mouse T helper epitope (RTEIHRNERTFT) and the flanking regions (**Fig. 1b**) but produced insufficient yields in the wheat-germ system for further study. The four other constructs were as follows: UF3 deleted the decoy peptide (peptide 1) and consisted of peptides 2-9; UF4 consisted of a 13 amino acid, non-natural, pan-DR-binding human T helper epitope [PADRE: AKFVAAWTLKAAA] and the UF3 sequence; the UF5 antigen was a dimer construct of UF3 + PADRE + UF3; and finally the UF6 antigen, a dimer construct similar to UF5 but connected with a glycine linker (GS(GGGS)_2_G) (**Fig. 1b, table S1**). These antigens were tested in immunization studies with a prime and two boost regimen (boosting on days 28 and 42) using outbred CD1 mice formulated with either Alhydrogel™ or GLA-LSQ adjuvant. The ranking criteria for progressing forward were (1) induction of an antibody response to both peptide 7 and 9, (2) induction of an antibody response to either peptide 7 or 9, and (3) functional T-B activity. Our initial immunological screen demonstrated that wheat germ expressed AnAPN1 constructs UF5 and UF6 were more immunogenic than the original AnAPN1 construct (UF1) (**fig. S2**). Based on serum endpoint titers alone, the immunogenicity of UF5 appeared to superior to that of other constructs. Antibodies generated using UF5:GLA-LSQ or UF6:GLA-LSQ were also found to be focused on the T-B epitope, peptide 9, and the recognition signals were significantly greater than that observed for UF1 (**fig. S2**).

Serum samples from the immunized mice were then used to further determine the T-B activity of the different antigens in both the standard membrane feeding assay (SMFA) using laboratory strains of *Plasmodium* (NF54) and by direct membrane feeding assay (DMFA) using circulating parasite strains in Cameroon (**fig. S3**). Including the wild-type *Plasmodium* strains helped us to further demonstrate the high T-B potential of the two “down-selected” antigen candidates (UF5 and UF6) that had also shown the highest endpoint titers in the immunization studies (**fig. S2**). However, our results indicated that there was no clear correlation between the antibody titer and T-B activity. Although we did not initially observe any advantage in terms of immunogenicity of formulating UF5 with GLA-LSQ over Alhydrogel™ (ALUM) (**fig. S2**), the subsequent functional studies suggest that the UF6:GLA-LSQ formulation produces the most effective T-B antibodies. Moreover, despite higher UF5:GLA-LSQ antibody titers, the UF6:GLA-LSQ antibodies proved to be more potent at blocking oocyst development in a DMFA (**fig. S3**).

### Establishing a large-scale CGMP production process for UF6b

Based on the finding that UF6 produced the most robust T-B antibodies, we developed a process to produce large scale CGMP quantities of the UF6 immunogen in preparation for First-In-Human clinical trials. In this process we made two changes to the antigen resulting in the “UF6b” immunogen: (*i*) removal of the C-terminal His-tag that was used for purification and (*ii*) addition of an inert, non-immunogenic peptide sequence, CGGSG at the C-terminus to protect from anticipated carboxyterminal protease clipping at higher fermentation scales (**table S1**). This construct was inserted into the *E. coli* expression vector pD1841 for expression under the control of phoA promoter in transformed BL21(DE3) cells (**Fig. 2)**. A Master Cell Bank was generated and used to produce UF6b (**Table 1**). The identified process produced 52.3 g of total UF6b at harvest (469.94 mg/mL in 111.3 L volume) in relatively low biomass of 45.8 g/L at the 120 L fermenter scale. This calculates to a specific productivity of 1% (0.010 g UF6b per g of *E. coli*). The pre-clinical grade bulk drug substance met all established acceptance criteria for quality attributes, including an endotoxin level of 6.7 EU/mg (**Table 2**).

**Fig. 2.**
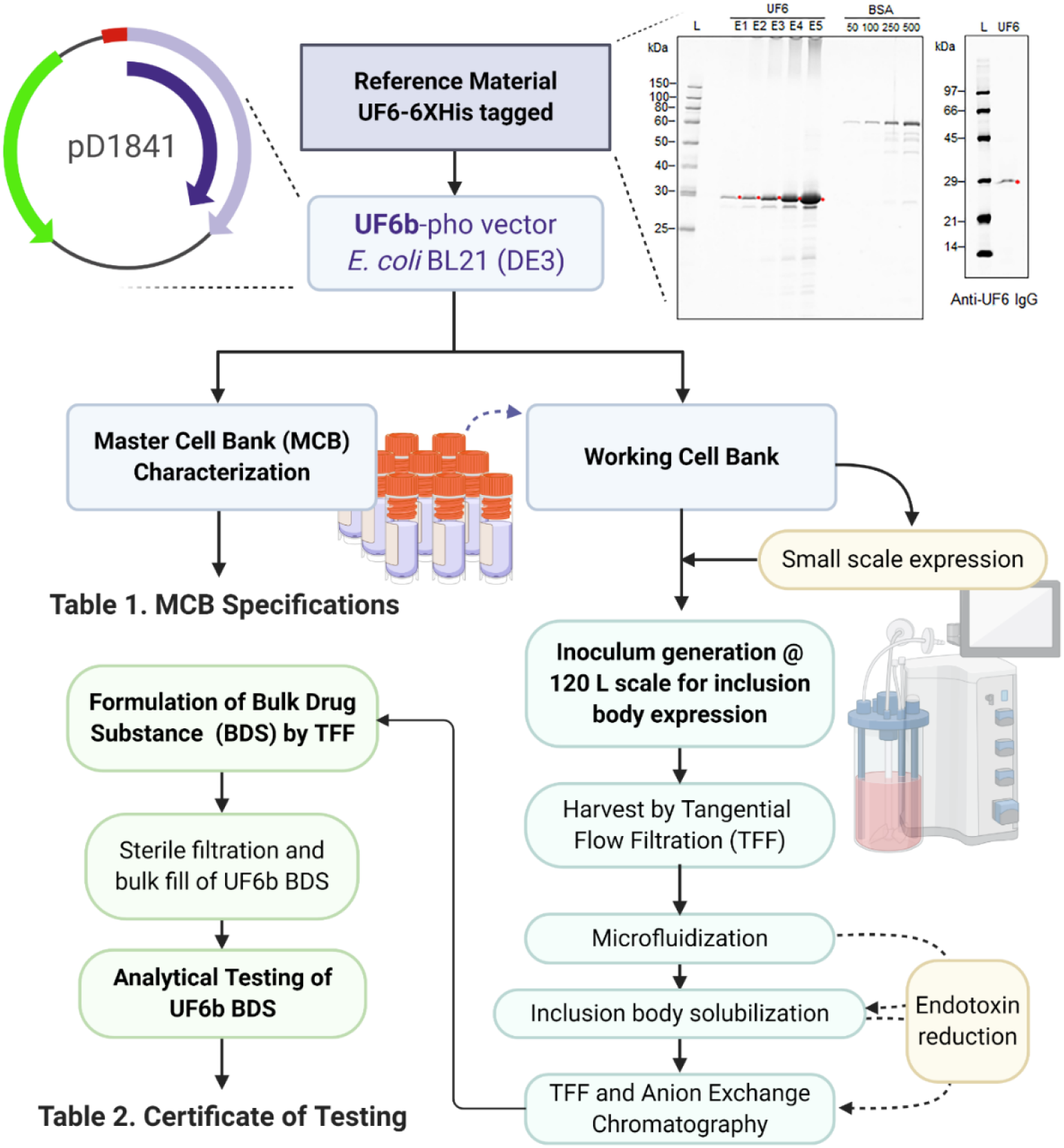
Process flow diagram for UF6b manufacture. (Top right) UF6 antigen produced by CellFree Sciences in the wheat germ cell-free system, which was recognized as a 29 kDa band by anti-UF6 antibody, was used as reference material by Ology Bioservices for process development. (Top left) *E.coli* BL21 (DE3) were transformed with UF6b-phopD1841 to produce a Master Cell Bank (MCB) expressing the UF6b, HIS-tag-free antigen. Using a lot from the MCB, pilot small-scale expression was followed by replicate 120-L scale runs with the goal of high yield production of UF6b in inclusion bodies to aid in the purification steps to reduce endotoxin. In-process UF6b was used for all immunological and functional assays. Bulk Drug Substance was further characterized and observed to yield low endotoxin levels. kDa, kilodalton. BSA, bovine serum albumin standards. See **Table 1** and **Table 2** for more details.

**Table 1.** UF6b-phoA-T7 Master Cell Bank (MCB)

| Analysis | Method | Acceptance Criteria | Results | Pass/Fail |
| --- | --- | --- | --- | --- |
| Viable Cell Count: Pre-freeze | TM-0088 | $\geq 1 \times 10^6$ CFU/mL | $3 \times 10^8$ CFU/mL | Pass |
| Viable Cell Count: Post-freeze | TM-0088 | $\geq 1 \times 10^6$ CFU/mL | $4 \times 10^8$ CFU/mL | Pass |
| Lysogenic bacteriophage | 510088GMP.BUK (BioReliance) | Bacteriophage not detected | Not detected | Pass |
| Mycoplasma | 300201GMP.BUK (BioReliance) | Negative for Mycoplasma | Negative | Pass |
| Purity testing of microbial cell banks | 510008GMP.BUK (BioReliance) | No evidence of contaminating organisms | Absent | Pass |
| Morphology | 510057GMP.BUK (BioReliance) | Gram negative, rod shaped bacterium | Conforms | Pass |
| Host Cell Phenotype | 512002GMP.BUK (BioReliance) | Profile consistent with host cell line | Conforms | Pass |
| Host Cell Genotype | 105058GMP.BUK (BioReliance) | Banding pattern of MCB consistent with host cell line | Conforms | Pass |
| Percentage of *E.coli* cells retaining expression vector | 104030GMP.BUK (BioReliance) | Report Results | 100% retain antibiotic resistance gene, 100% retain expression construct | N/A |
| Retention of the foreign UF6 gene: Sequencing | 106603GMP.BUK (BioReliance) | Conforms with the reference sequence for the coding sequence | Conforms | Pass |
| Plasmid Identity: Restriction Digestion | 104034GMP.BUK (BioReliance) | Coherent fragments with plasmid restriction map | Conforms | Pass |
| Retention of the recombinant construct – determination of plasmid copy number | 107315GMP.BUK (BioReliance) | Report Results | Test article AF85BF contained between 21.81 and 87.22 copies of AF85BF transgene target per cell (43.61 ± 2-fold) | N/A |
| Expression of ΔN-AnAPN1 (UF6) | TM-0087 | Anti-UF6b antisera identified a protein of MW 28 kDa | Conforms to Reference<sup>1</sup> | Pass |

Reference EVENT-2020-0039

**Table 2.** Certificate of Testing for UF6b Drug Product.

| Analysis | Method | Results |
| --- | --- | --- |
| pH | USP <791> | 8.05 |
| Visual Appearance | Visual Inspection | Free from visible particulates, slightly opalescence, slightly yellow |
| Identity | Western Blot | Conforms to Major Band in Reference Standard |
| Protein Concentration | Spectrophotometric/<br>A280 | 3.96 mg/mL |
| Purity | SEC-HPLC | Four peaks, in sequence high MW to low MW<br>Peak 1: 48.04%<br>Peak 2: 33.09%<br>Peak 3: 10.91%<br>Peak 4: 7.96% |
| Purity | RP-HPLC | 100% |
| Purity | CE-SDS-PAGE | 96.43% |
| Residual DNA | PicoGreen | 24.3 ng/mL |
| Residual HCP | ELISA | 206 ng/mg |
| Endotoxin | USP<1115> | 6.7 EU/mg |
| Bioburden | USP<85> | TAMC<sup>a</sup> = 0 cfu/mL<br>TYMC<sup>b</sup> = 0 cfu/mL |

<sup>a</sup>TAMC, Total Aerobic Microbial Count
<sup>b</sup>TYMC, Total combined yeasts and molds count

### UF6b is immunogenic and elicits anti-Peptide 9 antibodies in mice

To test the immunogenicity of the selected new preclinical UF6b product, outbred CD1 mice were immunized intramuscularly (20 μg/dose/mouse) following a prime plus two-boost regimen, with boosting on Days 28 and 70 to reflect an anticipated dosing interval in a clinical trial (**Fig. 3a, table S1**). UF6b alone (in the absence of adjuvant) is highly immunogenic in mice, achieving reciprocal serum endpoint titers of 10⁶ on Day 98 of the study (**Fig. 3b**), establishing its baseline immunogenicity. Mice were also immunized with UF6b formulated with either GLA-LSQ (**Fig. 3c**) or AddaVax™ (RUO version of the more potent MF-59 human-safe adjuvant to replace Alhydrogel™ as the adjuvanted UF6b positive control, **Fig. 3d**). The formulation of UF6b:GLA-LSQ elicited a higher, more rapid, and more stable response than either UF6b alone or UF6b:AddaVax™ (**Fig. 3e**).

**Fig. 3.**
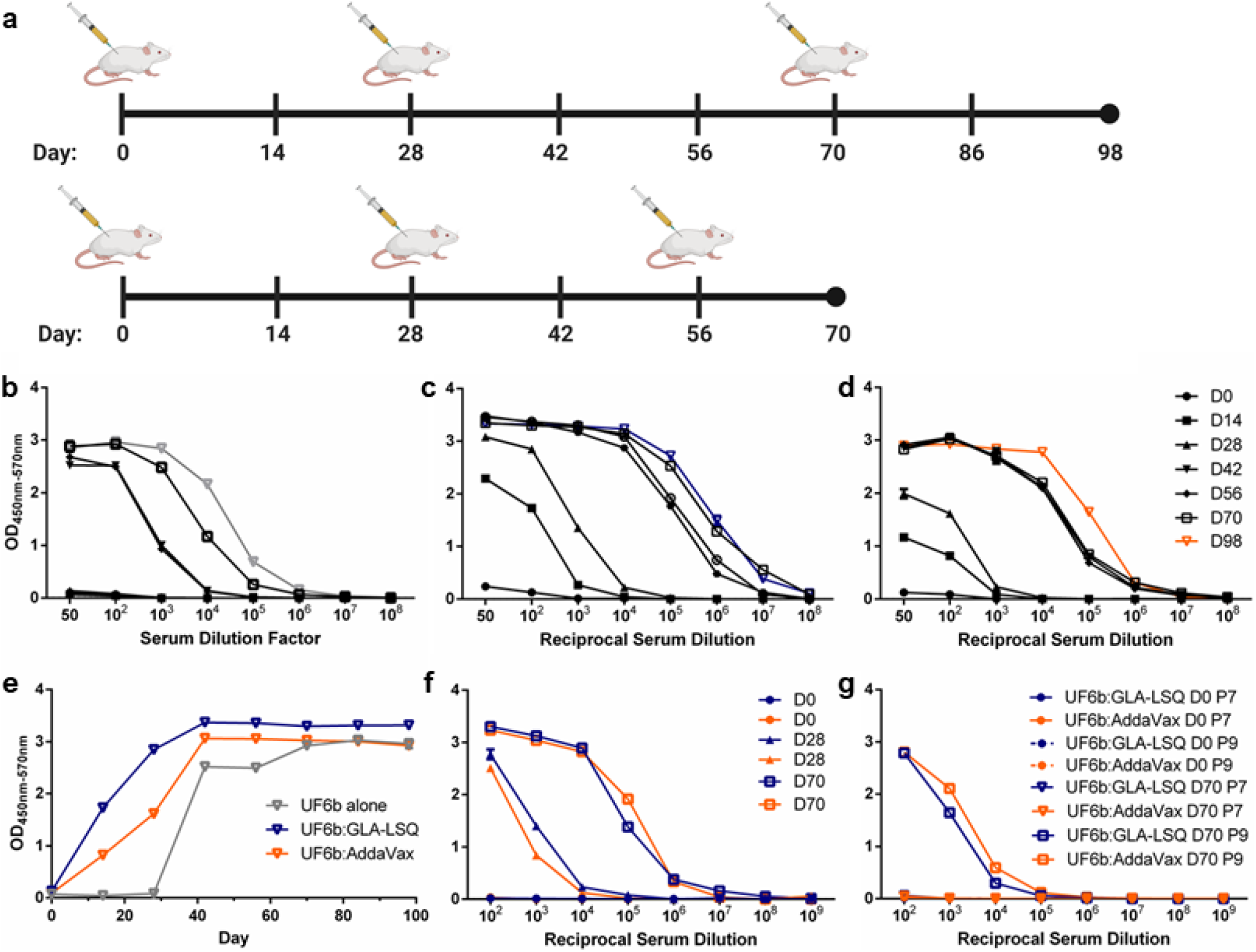
Immunogenicity of the UF6b construct. **(a)** Mice were immunized (i.m.) following a longer D98 prime and two-boost regimen (boosts on Days 28 and 70) or a compressed D70 prime and two-boost regimen (boosts on Days 28 and 56) (20 μg/dose/mouse). Analyses of the longer regimen sera are shown in **b-e**. **(b-d)** Mice were immunized with UF6b [grey] in the absence of adjuvant **(b)**, with UF6b:GLA-LSQ [blue] **(c)**, or with UF6b:AddaVax [orange] **(d).**(**e)** The kinetics of the antibody (1:100 dilution) response over time. Analyses of the shorter regimen sera are given in **f-g: (f)** UF6b given with either GLA-LSQ [blue] or AddaVax [orange] in the contracted D70 study. **(g)** ELISA analyses used pooled sera collected at the designated timepoints. Absorbances taken at 570 nm were deducted from absorbances at 450 nm to account for background. Error bars indicate SEM of triplicates. D, Day.

Considering that immunization regimens can influence the magnitude and durability of antibody responses to an immunogen, we also tested a Day 0 (prime) with a Day 28 and Day 56-boosting regimen to determine whether a more contracted 70-Day study would affect the overall humoral immune response in mice. We found that the contracted 70-Day immunization regimen (**Fig. 3a**) produced a comparable, antibody response to the 98-Day study with serum endpoint titers at ~1:10^8^ (**Fig. 3f**).

We hypothesized that deleting the N-terminal “decoy” peptide epitope (peptide 1) would focus the humoral response of immunized mice to the two key T-B epitopes, peptide 9 and peptide 7. We tested this hypothesis by conducting indirect peptide enzyme-linked immunosorbent assays (ELISAs) using only synthetic peptides for these two epitopes. Using sera collected from the 70-Day study as a representative humoral response for two different dosing regimens, we observed that the mice mounted an antibody response to peptide 9, with UF6b formulated with either adjuvant but failed to mount a response to peptide 7 (**Fig. 3g**). Both the longer and contracted studies were repeated with new cohorts of mice and the pooled antibody responses generated against UF6b and both key peptides remained similar (**fig. S4**). Importantly, a closer examination of individual mouse antibody titers from the 70-Day study indicated a high peptide 9 responder rate across all mice per group (**fig. S5-S6**).

### UF6b elicits potent T-B antibodies in mice

Previously, we have shown that immunizing mice with the N-terminal 125-aa recombinant AnAPN1 protein generates functional T-B antibodies^7–10^. In the current study, we assessed whether total IgG purified from mice immunized with the new construct UF6b similarly blocked parasite transmission by both SMFA and DMFA. The total IgG concentration of 50 μg/mL, corresponding to about 3.3 μg/mL of antigen-specific IgG, effectively blocked transmission of *P. falciparum* in *Anopheles gambiae* mosquitoes (**Fig. 4)**, leading to a significant reduction in oocyst intensity (i.e., the number of oocysts present per midgut). Antibodies raised in mice immunized with UF6b:GLA-LSQ were able to significantly reduce mean oocyst intensity greater than 90% (P<0.001, GLMM, Kruskal-Wallis, α<0.05) (**Fig. 4a, c**). Additionally, higher concentrations of total IgG did not appear to enhance T-B activity at concentrations above 50 μg/mL (**Fig. 4b**). This phenomenon has been described previously, wherein higher antibody concentrations have either resulted in plateauing of T-B activity or enhancement of parasite infection of the midgut; effects within the known intrinsic error of the assay^12–14^.

**Fig. 4.**
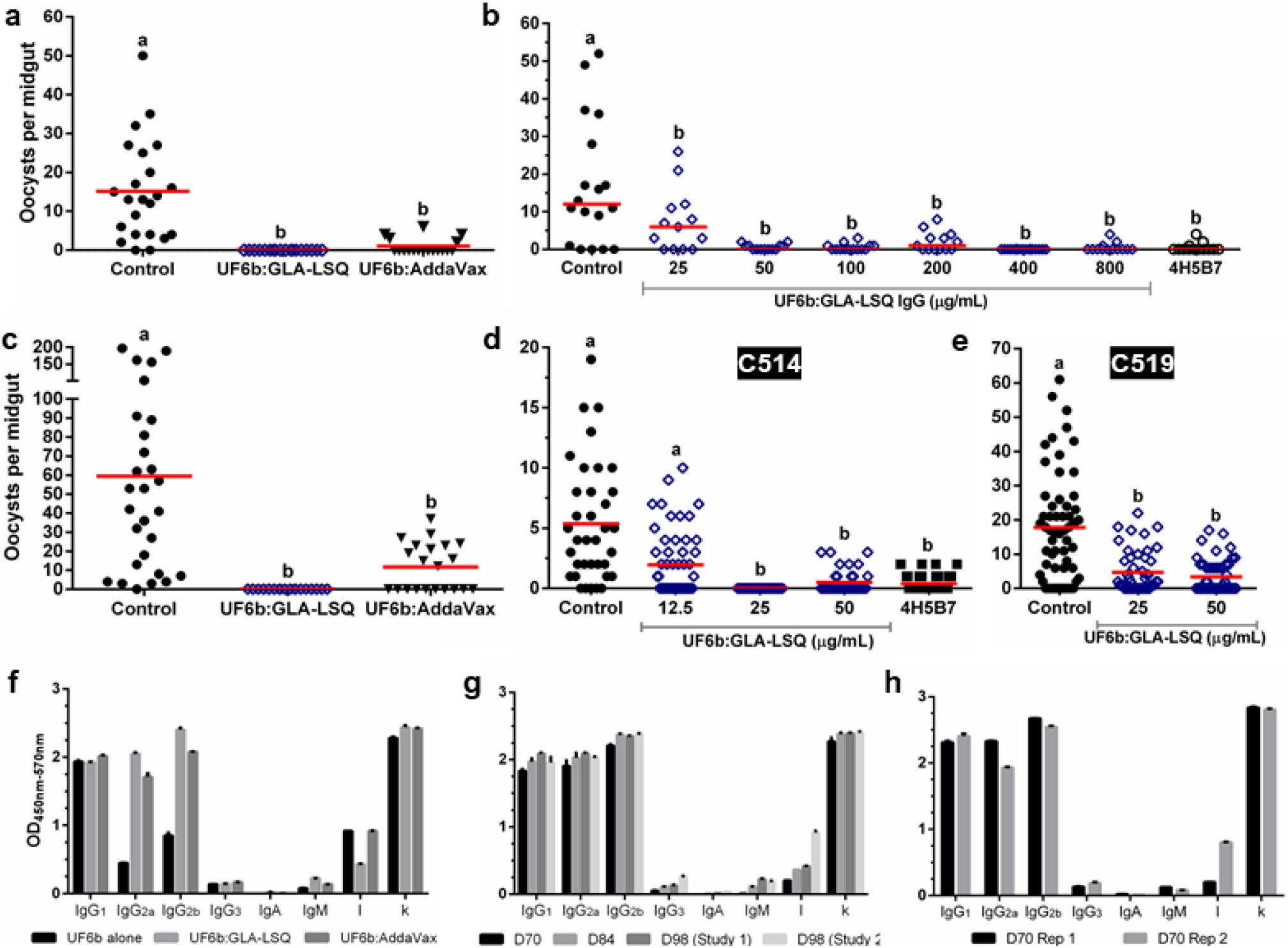
Functional activity of UF6b antibodies. (a-e) UF6b:GLA-LSQ generates potent transmission-blocking antibodies against both lab and naturally circulating *Plasmodium* strains. **(a)** Representative standard membrane feeding assay (SMFA) using 50 μg/mL total IgG purified from pooled sera of mice immunized with UF6b collected on Day 98. **(b)** Representative SMFA using serial dilutions of purified IgG from Day 70 of the D98 study. The monoclonal antibody (mAb) 4H5B7 was included at 250 μg/mL as a positive control. **(c)** Representative SMFA with a higher mean oocyst count in the control group, using 50 μg/mL total IgG purified from pooled sera of mice immunized with UF6b collected on Day 98. **(d-e)** Direct membrane feeding assay (DMFA) in Cameroon conducted with blood from 2 gametocytemic children (Case codes C514 and C519) with varying concentrations of total IgG. The peptide 7 mAb 4H5B7 served as the positive control. **(a-e)** Total IgG was purified from the same cohort of mice. Treatments without a common letter were found to be statistically significant (α = 0.05, p < 0.01, *N* =14-59) as calculated by Kruskal-Wallis test with Dunn’s post hoc correction or by zero-inflated Generalized Linear Mixed Model as appropriate. Red horizontal bar indicates mean oocyst number per mosquito midgut. **(f)** Comparison of Ig isotypes present in antisera between UF6b alone, UF6b:GLA-LSQ and UF6b:AddaVax™. **(g)** Comparison of Ig isotypes from collections at D70, D84, and D98 from the protracted dose regimen studies. **(h)** Ig isotypes for two replicates of the contracted dose regimen studies. **(f-h)** Sera used at 1:1,000 dilution. Error bars indicate the SEM for replicate wells.

We also tested the ability of purified total IgG from mice immunized with UF6b:GLA-LSQ to block naturally circulating *P. falciparum* parasites from infecting *An. gambiae* mosquitoes in Cameroon by DMFA. UF6b-specific IgG at 3.3 μg/mL resulted in reproducible reductions of 80-90% in mean oocyst intensity (P<0.001, GLMM, α<0.05) (**Fig. 4d,e)**.

To determine whether the observed differences in apparent T-B activity for antisera from UF6b:GLA-LSQ and UF6b:AddaVax immunizations can be explained by variances in antibody quality and quantity, we compared the immunoglobulin isotypes for these two groups as well as for UF6b alone and the contracted 70-Day dosing studies. In the absence of adjuvant, UF6b elicited primarily an IgG1 response with lower levels of IgG2a and IgG2b (**Fig. 4f**). When given in conjunction with either GLA-LSQ or AddaVax™, the immunoglobulin profile is more equally divided between these three subtypes. Also, between these three immunization groups no differences were noted for IgG3, IgA, and IgM, which were all comparably low and all groups had a higher proportion of kappa to lambda light chains (**Fig. 4f**). Very little variation in the immunoglobulin profile was seen in the D98 study after the D70 time point, although we did notice an increase in lambda light chain in the repeat D98 study (**Fig. 4g).** We noted that the contracted study of UF6b:GLA-LSQ produced a comparable immunoglobulin subclass profile (**Fig. 4h**) as compared to the longer D98 study (**Fig. 4g**), indicating flexibility in dose regimen schedules in future clinical studies. There was some variation in the amounts of IgG2a and lambda light chain between the two replicate 70-Day studies, which can be expected with outbred mice.

## Discussion

Here we successfully optimized our mosquito-based AnAPN1 candidate for further development and evaluation as a potent malaria TBV antigen. Moreover, our studies further demonstrated the benefits of using structural information in antigen optimization as a crucial step in the development of any malaria vaccine candidate. UF6b was reliably immunogenic in mice even when inoculated alone, unformulated with an adjuvant. The UF6b construct is based on a structure-function understanding of the immunogenic and mosquito-specific domains of the full-length APN1 from *An. gambiae*^10^. UF6b generated a peptide 9-specific response, which we have demonstrated previously^8^ confers potent T-B activity. While our original goal had been to target both peptides 9 and 7, which were predicted to strongly bind MHC II encoded by the HLA-DRB1 alleles common to East African populations^7,11^, we were unable to elicit a peptide response specific to peptide 7. This lack of a peptide 7-specific antibody response may result from an altered presentation of the peptide 7 epitope due to the flexibility of the glycine linker introduced in the new UF6b dimer construct. However, the lack of a peptide 7-specific response did not appear to affect the overall T-B activity of the antibodies, and achieves two priority goals: induction of antibodies to either peptide 7 or peptide 9 and demonstrated functional T-B activity at lower antibody concentrations.

In previous studies, between 0.8-1.6 mg/mL of purified total IgG from rabbits immunized with a near-full length recombinant AnAPN1 (i.e., UF1) was needed to block oocyst development in both SMFAs and DMFAs^7^. At this high total IgG concentration, it was estimated that only ~5-10 μg/mL were antigen-specific IgG. However, these studies used an immunogen containing the “immune decoy” epitope (peptide 1), causing a larger proportion of the murine IgG profile to target this malaria-irrelevant epitope^7,8^. We observed that a comparable amount of UF6b-specific IgG (3.3 μg/mL) was present in the antisera of mice immunized with UF6b:GLA-LSQ. These data suggest that the UF6b:GLA-LSQ formulation was indeed able to focus or skew the vertebrate immune response towards the peptide 9 epitope, resulting in a 32-fold lower concentration of total IgG (50 μg/mL as opposed to 1.6 mg/mL) needed to confer marked T-B activity. It remains to be seen if antibodies with similar potency can be induced in non-human primates or humans. Maximal T-B activity for total IgG following UF6b:GLA-LSQ immunization was comparable to that observed for the potent monoclonal antibody (mAb) 4H5B7, which targets peptide 7 only^8^. The degree of blocking activity appears to be related to the intensity of the infection of the mosquito in SMFAs and in DMFAs, the variation in T-B activity may be related to the genetic complexity of the gametocytes in the blood obtained from infected volunteers^15^. This is not entirely unexpected since the membrane feeder (although the gold standard for measuring T-B activity in the field) is artificial and circulation of infected red blood cells and antibody within the small feeders is not currently possible. As such, the distribution of antibody throughout the bloodmeal may not be even. The intrinsic error of membrane feeding assays notwithstanding^12–14^, complete blockade can be observed when oocyst intensities are less than 50 oocysts/midgut, which is a level expected in natural infections based on observations from field studies examining the range of possible oocyst intensities^16^.

Differences in T-B activity between the UF6b:GLA-LSQ and UF6b:AddaVax™ vaccine formulations cannot be explained by total antibody titers alone. Previous studies suggest that a predominantly Th2 response is elicited in mice following immunization with AnAPN1 (UF1) formulated with Alhydrogel™ ^[10]^. This is generally observed as lower IgG2 titers relative to IgG1 titers in immune sera. However, higher IgG2a and IgG2b titers to UF6b were observed with UF6b:GLA-LSQ and UF6b:AddaVax™ than UF6b alone, indicative of a Th1-biased humoral response in these outbred CD1 mice. UF6b:GLA-LSQ elicited a slightly higher IgG2a titer as compared to UF6b:AddaVax™ and more than a log higher IgG2a titer than UF6b alone. Saponin-based adjuvants (GLA-LSQ) and emulsions such as AddaVax™ or MF-59 are designed to induce robust and balanced Th1/Th2 responses, and this was clearly observed in our study for UF6b:GLA-LSQ and UF6b:AddaVax™ ^[17]^. The observation of a higher proportion of kappa light chains in antigen-specific IgG (all Ig subclasses) for the UF6b:GLA-LSQ has not been previously observed in studies using UF1:Alhydrogel™. In humans, variable light chains can influence antibody-binding specificity, serum half-life, and conformational flexibility in addition to other physicochemical properties, whereas subtle differences in these same properties have been described for variable heavy chains^18^. IgG3 responses have not been noted in our previous work with AnAPN1-targeted immunization, and although associated as a co-marker of a Th1 profile, the overall IgG3 titers in this study were low. Given that the potent mAb 4H5B7 is an IgG2b, kappa isotype, it is possible that the higher titers of IgG2a and IgG2b and light chain isotype ratio may have resulted in not only more antibody, but also a qualitatively better antibody repertoire. This hypothesis bears further study and could be relevantly and appropriately addressed in a Phase 1 clinical trial. It should be noted that mice are an imperfect model organism for predicting the success of malaria vaccines in humans, as mice generate different immune responses with distinct antibody characteristics as compared to non-human primates (NHPs) and humans^19^. As such, further testing of the tolerance, immunogenicity, and longevity of our UF6b:GLA-LSQ formulation are currently being completed in NHPs to allow a quicker transition to First-In-Human clinical trials.

The process development workflow (**Fig. 2**) established in this study positions the program well to move forward to CGMP production of UF6b. Based on a theoretical 0.1 mg/mL/subject as the highest dose in a Phase 1 study, the conservative estimated endotoxin level would be 0.35 EU/mL. This is already well below the <20 EU/mL recommendation for each dose of a recombinant protein vaccine and is 15-fold lower than the upper limit of endotoxin for Rubella vaccines^20^. In fact, the current process sets AnAPN1 (UF6b) endotoxin level within the reported range of several recommended pediatric vaccines.

Malaria TBVs are a critical component in the fight to eliminate and eradicate malaria. Our results meet and surpass the functional benchmark for a successful TBV, i.e., confer ≥80% T-B activity in terms of oocyst intensity and achieve this using lower concentrations of target-specific antibody titers in the membrane feeding assay^5^. Considering the inherent variation in vaccine response in human populations in malaria-endemic countries, partly as a result of acute malaria, concomitant infections and/or malnutrition^21^, it is important that the bar for the required concentration of antibodies with functional malaria T-B activity is lowered. Doing so would increase the likelihood of success that implementation of such a vaccine would result in demonstrable efficacy at the community level. The use of the GLA-LSQ adjuvant is predicted to help overcome limitations in the induced T follicular helper (Tfh) cell response, as circulating Tfh cells were found to be impaired in B cell help during an acute malaria episode^21^, and this adjuvant has proved to be critical in improving the total Tfh response in clinical studies with the Pfs25 transmission-blocking vaccine^22^. Overall, our results are promising and underscore the candidacy of the AnAPN1 molecule (exemplified by UF6b:GLA-LSQ) as a prime malaria TBV formulation. Coupled with the high-yield and low endotoxin level of the current pre-clinical Drug Product, the AnAPN1 TBV is positioned nicely to move forward with CGMP manufacture, lot release testing, and toxicity studies as a prelude to a Phase 1 clinical trial.

## Materials and Methods

### Experimental Design

For both SMFAs and DMFAs, IgG was purified from each 0.8-1 mL cardiac puncture sample per mouse. Since a single mouse cannot yield sufficient purified IgG for an assay, using current best practices, immune serum from either ten or twenty mice were pooled before purification. The purified IgG from the pooled immune sera permitted us to perform dose-response membrane feeding assays in triplicate.

Data collection was stopped at a predetermined timepoint of either Day 70 or 98. As per the animal care use committee approved protocol (protocol #201909359), data collection endpoints were also determined by the following humane endpoints for the test mice: if any of the test mice sustained greater than or equal to 15% weight loss from baseline weight or age-matched controls for those animals still maintained during the study; a body condition score of 2 or less; inability to reach food or water; impaired mobility; the presence of tumors; labored breathing, respiratory distress or cyanosis (blue tinged color); tremors, convulsions or seizures lasting more than 1 minute or occurring more than once a day; dehydration lasting over 24 hours that is unresponsive to treatment; or moribund - unable to right itself.

The objective of this study is to down-select to a new AnAPN1 construct capable of immunofocusing the humoral response to the two key T-B epitopes. Once this was accomplished, we sought to determine the immunogenicity and immunofocusing ability of the UF6b antigen as well as quantify the T-B activity of the elicited immune sera. To test if UF6b immunofocused the murine humoral response to peptides 7 and 9, mice were immunized via intramuscular injection (i.m.) with either UF6b alone, UF6b:GLA-LSQ, or UF6b:AddaVax™. At pre-specified timepoints blood samples were collected to perform ELISAs, SMFAs, and DMFAs. All data relevant data are included. The study was performed twice per dosing regimen with each mouse serving as a biological replicate in each cohort. Tests were performed with two-three technical replicates. There was no blinding in this study.

### Ethics

All research protocols have been approved by the relevant institutional review boards and national ethics committees for Cameroon (2016/03/730 and 2019/07/1168/CE/CNERSH/SP) and the University of Florida (UF) for concurrence and laboratory-based studies (IRB201800971, IRB201902261 and IRB201602150). Use of vertebrate animals (mice) for these studies have been approved by the UF Animal Care and Use Committee (UF Animal Welfare Assurance number A3377-01, Protocol# 201909359, approved on 06/07/2019).

### Wheat Germ Cell-Free Immunogen Expression

Templates for the expression of proteins UF1 to UF6 were prepared by gene synthesis (GenScript, USA). Before synthesis, sequences were optimized for expression in wheat, and a His-tag with six histidine residues was directly added without spacer to the C-terminus of the proteins. Templates were then cloned into the expression vector pEU-E01-MCS (CellFree Sciences, Japan) using the XhoI and BamHI restriction sites, and the DNA sequences of the templates confirmed before use in protein production. Plasmid DNA for each template was prepared using a Plasmid Maxi Kit (Qiagen, Germany). Protein expression was first confirmed by performing a 50 μL dialysis expression reaction using the wheat germ cell-free protein expression system extract WEPRO7240H (CellFree Sciences, Japan) according to the manufacturer’s directions for performing separate transcription and translation reactions. After a 72-h translation reaction, the crude reaction mixtures were centrifuged for 10 min at 21,500 x g to remove the insoluble fraction before loading onto a chelated-Ni resin (Ni-NTA Agarose, Qiagen, Germany). Proteins were purified on the Ni resin using a commercial buffer set (His Buffer Kit, GE Healthcare, USA). In brief, crude reaction mixtures were centrifuged at 10 min at 21,500 x g to separate the soluble from the insoluble fraction. The samples were diluted 4-fold using a buffer containing 20 mM Na-phosphate, 0.5 M NaCl, 0.02 M imidazole (pH 7.5) before loading onto the resin. After the binding reaction, the resins were washed three times with a 20 mM Na-phosphate, 0.5 M NaCl, 0.05 M imidazole buffer before the proteins were eluted twice with 20 mM Na-phosphate, 0.5 M NaCl, 0.5 M imidazole Elution Buffer. Proteins from each fraction during expression and later processing were resolved by 15% SDS-PAGE (DRC, Japan) and stained with Coomassie Brilliant Blue. Different amounts of BSA were run in parallel to quantitate yields of the expressed proteins. Because protein UF2 was partially insoluble and expressed at lower yields compared to the UF1 protein, further studies with UF2 expression were discontinued in favor of UF1. Remaining proteins UF1 and UF3 to UF6 were produced on a larger scale using 3 mL dialysis translation reactions following the same procedures established during the expression tests. As a part of large-scale protein preparation, a buffer containing 15% sucrose, 10 mM Tris, 0.2% Tween-80 (pH 7.8) replaced the Elution Buffer. Proteins were shock-frozen in liquid nitrogen and stored at −80°C until later use.

### Process Development of UF6b Manufacture

UF6 sequence was modified to remove C-terminus His-tag and in its place, we added a protective C-terminal, inert peptide linker sequence (CGGSG) to generate UF6b^23^. The produced sequence (GenScript) was cloned into the expression vector pD1841 under phoA promoter control and kanamycin as a selection marker. Competent *E. coli* BL21(DE3) (ThermoFisher) were transformed and screened for expression of UF6b by Western blot. Research cell banks (RCB) and Master cell bank (MCB) were generated for production of UF6b. Process development was performed in a scale-down models in a 10 L glass-vessel bioreactor (BioFlo 320, Eppendorf). Established process parameters were then scaled up to 120 L in a BioFlo Pro 150 stainless steel fermenter (Eppendorf). Phosphate depletion during the growth phase induced UF6b expression into inclusion bodies. Harvest was performed via tangential flow filtration using 750 kDa hollow fiber cartridge (Spectrum/Repligen) at load challenge <50 g/m^2^. The biomass was concentrated to >100 g/L and buffer exchanged into 50 mM HEPES (pH 8.0) buffer and subsequently diluted to 50 g/L for mechanical lysis by microfluidization (Microfluidics M-110E) at 20,000 psi for four (4) cycles with temperature controlled ≤16 °C. The insoluble inclusion body (IB) fraction was separated by centrifugation at 10,000 x g for 30 minutes at 2-8 °C (Sorvall Lynx 6000, ThermoFisher). The IB pellet was washed with 20 mM HEPES (pH 8.0), 2 M urea, 1% Triton X-100 at a 1:10 ratio (g of WCW biomass to wash buffer) and isolated by centrifugation. Two-phase separation wash with Triton X-114 was performed to reduce endotoxin levels^24^. The washed pellet was then solubilized with 6 M urea in 20 mM HEPES (pH 8.0) buffer for 1 h with mild agitation and an additional endotoxin removal step performed on the soluble preparation as previously referenced. Solubilized UF6b was refolded by dilution (from 6 M urea to 1 M urea) with 20 mM HEPES (pH 8.0) and buffer exchanged into 20 mM Tris-HCl (pH 8.0) + 15% sucrose + 200 mM NaCl by TFF to prepare the material for further purification by anion exchange chromatography (AEX).

UF6b polishing was performed by tandem AEX on Capto Q ImpRes (Cytiva) medium at linear velocity of 150 cm/h in flow-through mode followed by bind and elute mode. The effluent from flow-through mode was diluted with 20 mM Tris-HCl (pH 8.0) to achieve conductivity <7 mS/cm. After equilibration with 20 mM Tris-HCl (pH 8.0), binding of UF6b to AEX at ≤50 mg/mL resin was followed by a 5-column volume (CV) wash with equilibration buffer and eluted by linear gradient from 0 to 500 mM NaCl over 15 CV. Peak fractions were further analyzed for field and endotoxin content and pooled. Formulation was performed by TFF (ProStream mPES,1 m^2^ 10 kDa MWCO, L-screen; Repligen) operated with a TMP of 15 psig at a process flux of 8 LMH. The retentate was concentrated to >1 mg/mL then buffer exchanged with 7 diavolumes of 20 mM HEPES, 15% sucrose, 0.2% Tween 20 (pH 8.0). After the retentate was recovered, Tween-20 was added to Bulk Drug Substance (BDS) to a final concentration of 0.2%, and the BDS was filtered through a sterilizing grade 0.2 μm filter (Opticap PES, Millipore Sigma) to generate formulated BDS. BDS was aliquoted into HDPE bottles and stored at ≤65 °C.

### Mouse Immunizations

To down-select to the optimal AnAPN1 construct, female outbred CD1 mice (N=5) aged 5-7 weeks (Charles River) were immunized with 20 μg of either wheat germ cell-free expressed UF1, UF3, UF4, UF5, or UF6 emulsified 1:1 with GLA-LSQ or Alhydrogel. Mice were primed subcutaneously and boosted on Days 28 and 42 intraperitoneally (i.p.) for antigens formulated with Alhydrogel and intramuscular (i.m.) for antigens formulated with GLA-LSQ. Sera was collected on Day 0 and prior to boosting on Days 28 and 42. On Day 56 the mice were sacrificed, and sera collected via cardiac puncture.

To test the new UF6b construct, female CD1 outbred mice (Charles River) aged 6-8 weeks were used. Forty mice were split into three groups (**table S1**): Group A contained 10 mice immunized (i.m.) with UF6b; Group B (main focus group) contained 20 mice immunized (i.m.) with UF6b:GLA-LSQ; and Group C contained 10 mice immunized (i.m.) with UF6b:AddaVax™. For all treatment groups, mice received 20 μg UF6b in diluent (20 mM HEPES, 15% sucrose, 2% Tween-20, pH 8.0) to 100 μL total volume without or with adjuvant. For treatment groups B and C described above, 50 μL adjuvant was mixed 1:1 (vol/vol) with 50 μL antigen in diluent (total volume = 100 μL). At Day 0, mice received an i.m. injection of 50 μL of the antigen/adjuvant formulation in each caudal thigh, resulting in a total priming dose volume of 100 μL. At Days 28 and 70 post-priming, all mice were boosted i.m. with the same formulations described above (**Fig. 3a**). Every two weeks post-priming, mice were bled to collect sera for anti-AnAPN1 antibody titer determination via ELISA. Mice were sacrificed four weeks after the final boost and serum was collected via cardiac puncture. A single replicate study was completed with 10 mice immunized with the same UF6b:GLA-LSQ formulation in the same manner and timepoints. Unfortunately, due to COVID-19 closures, Day 84 sera could not be collected.

Two replicate studies with a contracted dosing schedule were completed with 20 female CD1 mice (Charles River) and used at 6 to 8 weeks of age. Ten mice were immunized with the same UF6b:GLA-LSQ formulation described above, and ten mice received the UF6b:AddaVax™ formulation. All mice received a prime dose i.m. on day 0 and were subsequently boosted on Days 28 and 56 post priming (**Fig. 3a**). Sera was collected every two weeks post-priming until the endpoint at Day 70 when the mice were sacrificed.

### Indirect Enzyme-Linked Immunosorbent Assays

Maxisorp 96-well ELISA plates (Nunc, Fisher Scientific) were incubated overnight at 4°C with 1 μg/mL antigen in 0.1 M PBS (pH 7.2). After three washes with PBS-Tween 20 (0.1%) (PBST20), the plates were blocked for 1 h at room temperature with 1% bovine serum albumin (BSA) in 1xPBS. Serum samples were diluted to 1:50 and 1:100, then diluted ten-fold to 1:10^8^ in 0.5% BSA in 1xPBS. After discarding the BSA and drying the plates, 100 μL of each of the eight serum dilutions were added to each well in triplicate and incubated 1 h at room temperature. Plates were again washed three times with PBST20. For detection, 100 μL of a horseradish peroxidase (HRP)-conjugated goat anti-mouse IgG(H+L) (KPL) diluted 1:5,000 in 0.5% BSA was added to each well and incubated for 1 h at room temperature. Plates were washed three times with PBST20 and developed by adding 100 μL of (3,3′,5,5′-tetramethylbenzidine) microwell peroxidase substrate (KPL) to each well. Development was stopped after 5 min by the addition of 100 μL of 1 M phosphoric acid. A Synergy™ HTX Multi-Mode Microplate Reader was used to read the OD values at 450 nm and 570 nm. The OD value at 570 nm was subtracted from the OD value at 450 nm to correct for background. Serum end point titers were defined as the reciprocal serum dilution giving a higher density reading than the average OD value at 450 nm-570 nm of the pre-immune serum plus three standard deviations.

### Peptide Indirect Enzyme-Linked Immunosorbent Assays

Maxisorp 96-well ELISA plates (Nunc, Fisher Scientific) were coated with 100 μL of Poly-L-Lysine (PLL) at a concentration of 50 μg/mL in 0.05 M sodium bicarbonate buffer (pH 9.6) and incubated covered for one h at room temperature. The plates were washed once with PBST20, then coated in 100 μL 1% (v/v) glutaraldehyde in PBS and incubated covered for 15 minutes at room temperature. The plates were washed again once with PBST20, and then coated with 100 μL of either peptide 1, peptide 7, or peptide 9 *(7,8)* and incubated covered overnight at 4°C. The next day the plates were washed twice with PBST20, then 200 μL 1M glycine was added to each well and plates were incubated covered for one h. The plates were washed twice with PBST20 and 300 μL of a 1:1 solution of 5% milk and 1% gelatin was added to the plates and incubated covered for one h. The serum samples were diluted in PBS using serial ten-fold dilutions from 1:10^2^ to 1:10^9^. After washing the plates once with PBST20, each of the eight sample dilutions was added to each well in triplicate and incubated covered for one h. The plates were washed three times with PBST20 and 100 μL (HRP)-conjugated goat anti-mouse IgG(H+L) (KPL) diluted 1:5000 in PBS was added to each well and incubated covered for one h at room temperature. Plates were washed three times with PBST20 and developed by adding 100 μL of (3,3′,5,5′-tetramethylbenzidine) microwell peroxidase substrate (KPL) to each well. Development was stopped after five min by the addition of 100 μL of 1 M phosphoric acid. A Synergy™ HTX Multi-Mode Microplate Reader was used to read the OD values at 450 nm and 570 nm. The OD value at 570 nm was subtracted from the OD value at 450 nm to correct for background. Serum end point titers were defined as the reciprocal serum dilution giving a higher density reading than the average OD value at 450 nm-570 nm of the pre-immune serum plus three standard deviations.

### Mosquito colony

*An. gambiae* (Keele) was used in all experiments. Mosquitoes were reared under standard insectary conditions of 27°C, 80% relative humidity, and 12 h light:12 h dark cycle. Eggs were hatched in milli-Q water supplemented with 0.02% yeast slurry. The emerging larvae were reared in plastic trays (25 cm long × 20 cm wide × 14 cm high) at a density of 300-400 larvae per tray and provided a daily ration of 1g koi fish food per tray/day. Pupae were collected and put in holding cages for emergence. Emerged adults were fed *ad libtum* on 10% sucrose solution. For all experiments, 5-to-6-day old adult females were used.

### Standard Membrane Feeding Assays

*P. falciparum* (NF54) gametocyte cultures were produced as previously described^25^. Briefly, cultures were seeded at 0.5% parasitemia and grown under hypoxic conditions in RPMI culture media supplemented with HEPES, hypoxanthine, glutamine, and 10% heat-inactivated human serum. The RPMI media was changed daily during culturing. Gametocyte cultures were harvested 17-18 days after initiation, and infected red blood cells were brought up in heat-inactivated human serum (LifeSouth Community Blood Center Inc, Gainesville, FL) plus human RBCs (blood type O-positive) at 1% gametocytemia and 30% hematocrit. Infective blood was mixed with control (IgG purified from naïve mouse serum) or total IgG purified from pooled sera of mice immunized with UF6b prior to delivery directly into water-jacketed membrane feeders maintained at 37°C via a circulating water bath. The final concentration of total IgG in 300 μL total volume of infective blood was 50 μg/mL for each technical replicate. Female *An. gambiae* were starved for 6 h, placed in paper cups with tops covered with netting (n = 40 per cup) and allowed to feed from each feeder for 20 min. Fully engorged female mosquitoes were maintained for 8 d when they were dissected for midgut oocyst enumeration. The dissected midguts were stained with 0.1% mercurochrome to visualize oocysts. A total of three biological replicate experiments were conducted.

### Direct Membrane Feeding Assays

Mosquito infections were conducted during the rainy seasons in 2019. Procedures for gametocyte carrier detection, blood collection and mosquito infections were performed as previously described^24^. Briefly, *P. falciparum* gametocyte carriers were identified among asymptomatic children (ages 5 to 11) in primary schools in the Mfou district, 30 km from Yaoundé, Cameroon. Venous blood was drawn from gametocyte-positive individuals in heparinized Vacutainer tubes from the antecubital fossa. Membrane feedings were set using donor’s blood with replacement of the serum by a non-immune AB serum. The local laboratory strain of *An. coluzzii*, named Ngousso, was used for mosquito feedings. Following infection, mosquitoes were maintained at the laboratory under standard insectary conditions (27 ± 2°C, 85 ± 5% RH, 12h light/dark) for 8 days until dissections.

### IgG purification and antigen-specific IgG quantification

Total IgG was purified from pooled serum using Protein A/G Magnetic Agarose Beads (Thermo Fisher Scientific) according to manufacturer’s instructions. The antibodies were quantified using Peirce™ BCA protein assay kit. Antigen-specific IgG was purified from pooled serum with UF6b-conjugated Dynabeads™ M-270 Epoxy (Thermo Fisher Scientific) following manufacturer’s instructions. The antibody titer of the purified UF6b-specific IgG and total IgG (serum) was determined with the Easy-Titer™ Mouse IgG assay kit (Thermo Fisher Scientific).

### Antibody Isotyping

Anti-UF6b immunoglobulin isotypes were determined with the Mouse Typer Isotyping Panel (Bio-Rad). Plates were coated and blocked, as described above, and mouse sera diluted 1:1,000 was added. After washing five times the isotyping panel was added, followed by five washes, then (HRP)-conjugated goat anti-rabbit IgG(H+L) (KPL). Absorbances were detected as described above.

### Statistical Analysis

To determine differences in oocyst intensity (defined by either the mean oocyst count) between pre-immune sera and test sera in SMFAs the data was first cleaned of outliers using a robust nonlinear regression test (ROUT) and a Q coefficient of 1% for all treatment and control groups^27^ using GraphPad Prism 6. After comparing the raw and cleaned data and considering that assumptions and overall significance results were not affected, outliers were removed to account for the intrinsic noise in the feeding assays, contributed in part by the uneven distribution of spiked antibody and pooling of gametocytes in the membrane feeder as well as variability in mosquito feeding voracity. The cleaned data was then analyzed by either a zero-inflated or normal Generalized Linear Mixed Model statistical test (GLMM) using the *R* package^28^, and when neither was appropriate, a non-parametric Kruskal-Wallis test followed by Dunn’s post hoc test using GraphPad Prism 6^7,8,10,29^.

## Data availability

All materials generated through this study are available following a request with the corresponding author.

## Acknowledgments

**General**: The authors thank Drs. Hilary Hurd and Paul Eggleston for the *An. gambiae* (Keele) strain and Paul Howell for initial assistance with mosquito colony maintenance. The authors thank especially the children, parents, and teachers in Mfou, Cameroon as well as other staff members of the Centre Pasteur du Cameroun for their involvement in and contributions to this study. Figures 1 and 2 were created in BioRender.

**Funding:** This work was funded by the Global Health Innovative Technology Fund (grant numbers G2015-214 and G2017-211).

## Author contributions

R.D., M.H. designed and directed the overall project. N.B., P.K., R.E.T. and V.N. performed experiments and analyzed data. V.N. and D.M. maintained the mosquito colony. B.L.G., T.H., and A.T. maintained the parasite cultures. J.M. E.V., R.C., directed and conducted process development activities for the UF6b product. M.H. directed the production of initial immunogen constructs and reference standards. R.F.H. provided GLA-LSQ and SE adjuvants and immunogenicity study design recommendations. S.N. conducted membrane-feeding studies in Cameroon. N.B., M.H., J.M. and R.D. wrote the draft and final manuscript with support from all authors.

