## Supplementary Materials for "Immunofocusing humoral immunity potentiates the functional efficacy of the AnAPN1 malaria transmission-blocking vaccine antigen"

### Supplementary Table S1. AnAPN1 v2.0 immunogen constructs.

| Construct / Amino Acid Sequence | pI / MW (Da) |
| --- | --- |
| **Construct UF1:**<br>MCDERYRLPTTSIPIHYDLHLRTEIHRNERTFTGTVGIQLQVVQATDKLVMHNRGLVMSS<br>AKVSSLPNGVTGAPTLIGDVQYSTDTTFEHITFTSPTILQPGTYLLEVAFQGRLATNDDG<br>FYVSSYVADNGERRYLAHHHHHH | 6.39 / 16,105.05 |
| **Construct UF2:**<br>MCIQLQVVQATDKLVMHNRGLVMSSAKVSSLPNGVTGAPTLIGDVQYSTDTTFEHITFT<br>SPTILQPGTYLLEVAFQGRLATNDDGFYVSSYVADNGERRYLAHHHHHH | 6.19 / 11,927.40 |
| **Construct UF3:**<br>MCDLHLRTEIHRNERTFTGTVGIQLQVVQATDKLVMHNRGLVMSSAKVSSLPNGVTGA<br>PTLIGDVQYSTDTTFEHITFTSPTILQPGTYLLEVAFQGRLATNDDGFYVSSYVADNGER<br>RYLAHHHHHH | 6.33 / 14,261.98 |
| **Construct UF4:**<br>MCAKFVAAWTLKAAADLHLRTEIHRNERTFTGTVGIQLQVVQATDKLVMHNRGLVMSS<br>AKVSSLPNGVTGAPTLIGDVQYSTDTTFEHITFTSPTILQPGTYLLEVAFQGRLATNDDG<br>FYVSSYVADNGERRYLAHHHHHH | 6.79 / 15,591.58 |
| **Construct UF5:**<br>MCDLHLRTEIHRNERTFTGTVGIQLQVVQATDKLVMHNRGLVMSSAKVSSLPNGVTGA<br>PTLIGDVQYSTDTTFEHITFTSPTILQPGTYLLEVAFQGRLATNDDGFYVSSYVADNGER<br>RYLAAKFVAAWTLKAAADLHLRTEIHRNERTFTGTVGIQLQVVQATDKLVMHNRGLVMS<br>SAKVSSLPNGVTGAPTLIGDVQYSTDTTFEHITFTSPTILQPGTYLLEVAFQGRLATNDD<br>GFYVSSYVADNGERRYLAHHHHHH<br><br>AKFVAAWTLKAAA (PADRE Linker) | 6.38 / 28,778.37 |
| **Construct UF6:**<br>MCDLHLRTEIHRNERTFTGTVGIQLQVVQATDKLVMHNRGLVMSSAKVSSLPNGVTGA<br>PTLIGDVQYSTDTTFEHITFTSPTILQPGTYLLEVAFQGRLATNDDGFYVSSYVADNGER<br>RYLAGSGGGGSGGGGSGGGGGSGDLHLRTEIHRNERTFTGTVGIQLQVVQATDKLVM<br>HNRGLVMSSAKVSSLPNGVTGAPTLIGDVQYSTDTTFEHITFTSPTILQPGTYLLEVAFQ<br>GRLATNDDGFYVSSYVADNGERRYLACGGSGHHHHHH<br><br>GS[GGGGS]$_2$GGGGSG (Glycine Linker) | 6.13 / 29,014.22 |
| **Construct UF6b:**<br>MCDLHLRTEIHRNERTFTGTVGIQLQVVQATDKLVMHNRGLVMSSAKVSSLPNGVTGA<br>PTLIGDVQYSTDTTFEHITFTSPTILQPGTYLLEVAFQGRLATNDDGFYVSSYVADNGER<br>RYLAGSGGGGSGGGGSGGGGGSGDLHLRTEIHRNERTFTGTVGIQLQVVQATDKLVM<br>HNRGLVMSSAKVSSLPNGVTGAPTLIGDVQYSTDTTFEHITFTSPTILQPGTYLLEVAFQ<br>GRLATNDDGFYVSSYVADNGERRYLACGGSG<br><br>GS[GGGGS]$_2$GGGGSG (Glycine Linker) | 5.71 / 28,191.37 |

Calculated by https://web.expasy.org/compute_pi/

**Table S2. Immunization groups.**

| Experimental Groups | Dose | Number of mice/groups | Volume of UF6b-stock (0.4 mg/mL) | Volume of Adjuvant |
|---|---|---|---|---|
| **A**: UF6b alone | 20 µg | 10 | 0.6 mL | (0.6 mL diluent buffer instead of adjuvant) |
| **B**: UF6b:GLA-LSQ | 20 µg | 20 | 1.2 mL | 1.2 mL |
| **C**: UF6b:AddaVax™ | 20 µg | 10 | 0.6 mL | 0.6 mL |

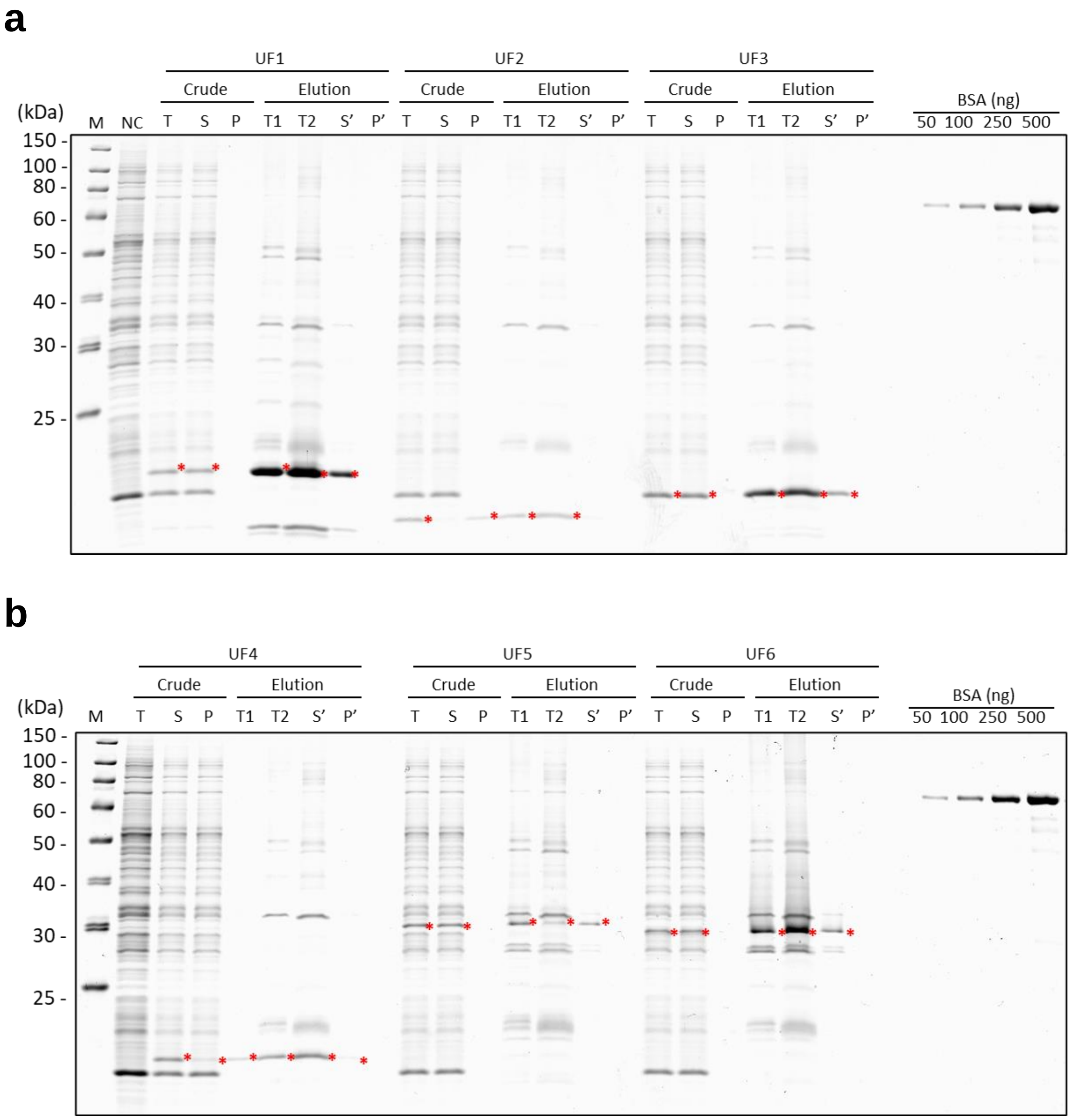


**Fig. S1. Expression and purification test for proteins UF1 to UF6 using a wheat germ cell-free protein expression system.** Proteins UF1-UF3 **(a)** and UF4-UF6 **(b)** were expressed for 72 h in 50 µL translation reactions using a wheat germ cell-free protein expression system followed by purification on a Ni resin using a His-tag at the C-terminus of the proteins and isolated after centrifugation as further outlined in the Methods. All fractions were analyzed on 15% SDS-PAGE along with a BSA standard: M = Molecular weight marker as indicated in the figures; T = Total fraction from crude translation reaction mixture; S = Soluble fraction taken from crude translation reaction mixture after centrifugation; P = Pellet fraction taken from crude translation reaction mixture after centrifugation; T1 = Elution fraction 1; Elution fraction 2; S' = Soluble fraction of elution fractions after centrifugation; P' = Pellet fraction taken from elution fractions after centrifugation; BSA = reference standard; * = indicating expressed proteins.

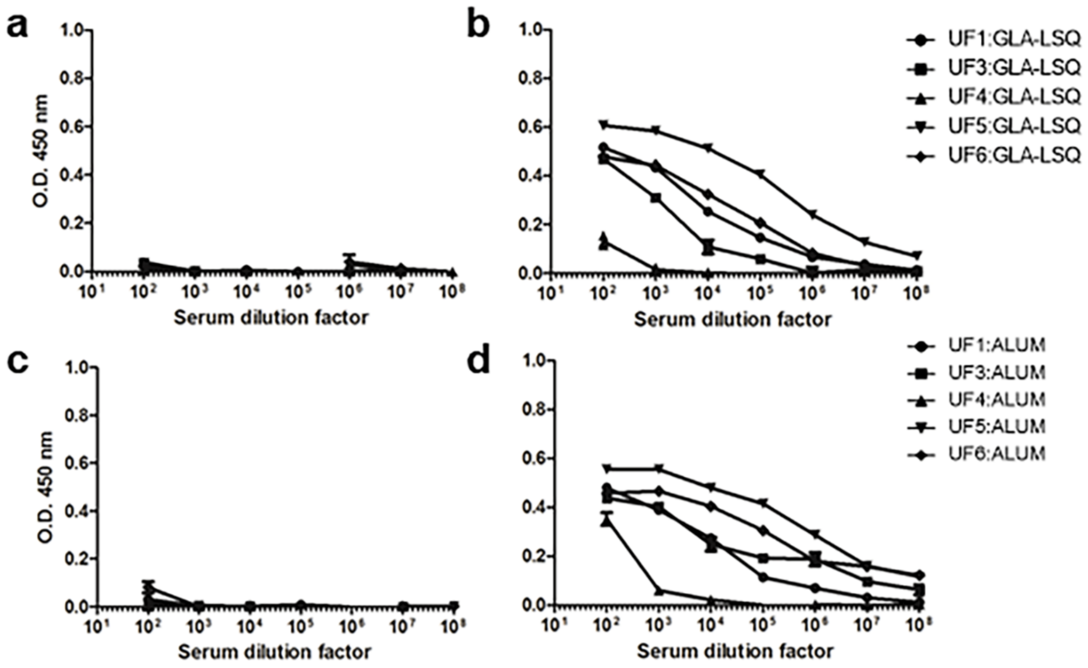


**Fig S2. Initial immunogenicity screen of GLA-LSQ and Alhydrogel™ formulations with UF3, UF4, UF5 or UF6 as compared to UF1. (a)** Pre-immune sera (N=5 mice) for wheat germ cell-free expressed antigens formulated with GLA-LSQ adjuvant. **(b)** Day 70 immune sera for antigens formulated with GLA-LSQ adjuvant. **(c)** Pre-immune sera (N=5 mice) for antigens formulated with

Alhydrogel (ALUM) adjuvant. **(d)** Day 70 immune sera for antigens formulated with Alhydrogel™ (ALUM) adjuvant. Error bars indicate SEM of triplicates.

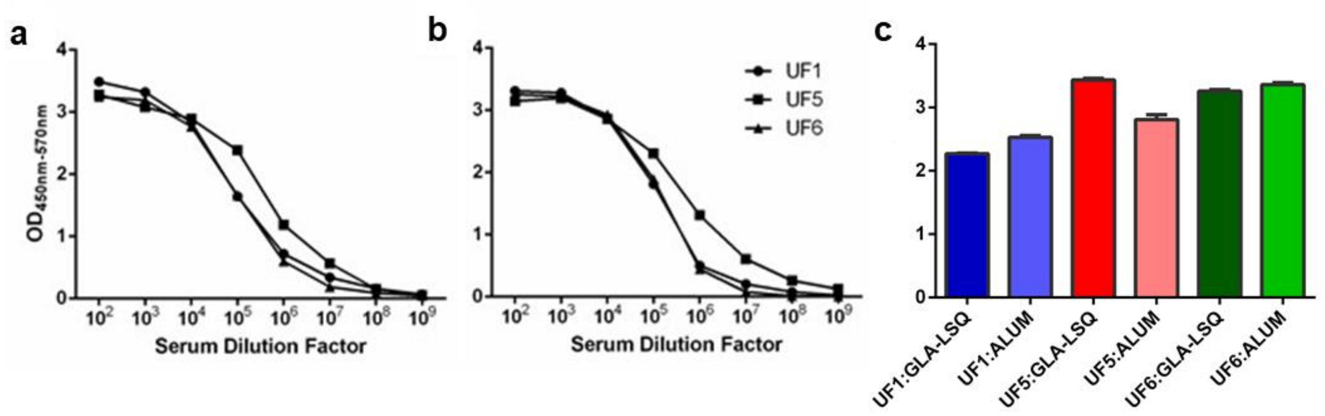


**Fig. S3. Comparison of GLA-LSQ and Alhydrogel™ formulations with UF1, UF5 and UF6. (a-b)** Anti-UF1, UF5, and UF6 antibody titers from mice immunized with 20 µg/mL of antigen were determined by indirect ELISA. The antigen was formulated with GLA-LSQ (**A)** or Alhydrogel**™** (indicated as ALUM) (**b**). **(c)** Anti-Peptide 9 titers were determined by indirect peptide ELISA with sera diluted 1/1,000. Error bars indicate SEM of triplicates.

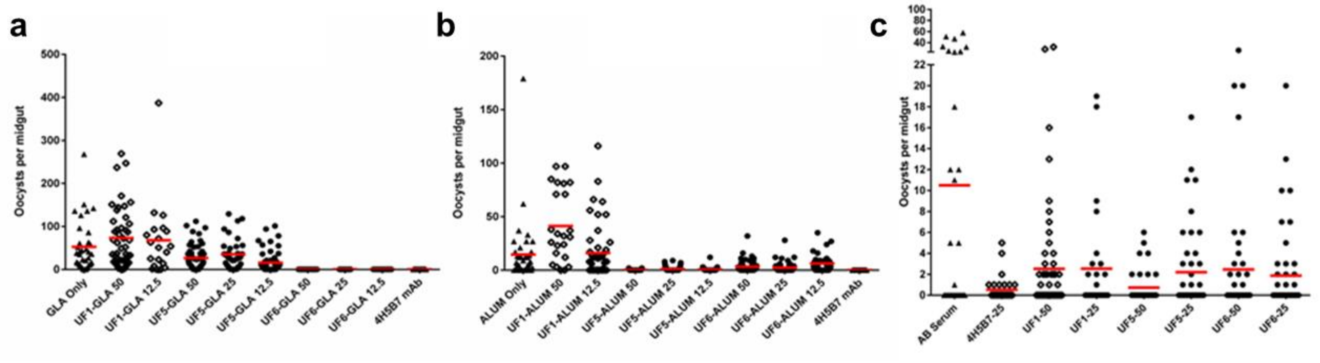


**Fig. S4. Screening Study: Functional transmission-activity assessment.** Direct Membrane Feeding Assay (DMFA) conducted with UF1, UF5, and UF6 either with GLA-LSQ **(a)** or Alhydrogel™ (indicated as ALUM) **(b)** performed using infectious, gametocytemic blood from a single carrier during the May-July transmission season in Cameroon. Antibody concentrations (ranging from 12.5-50 µg/mL) per feeder are indicated. **(c)** Representative SMFA with UF1, UF5, and UF6 formulated with GLA-LSQ. 4H5B7 is a monoclonal antibody (mAb) to peptide 7, which serves as a positive control. In panel A-B, 4H5B7 was used at a single concentration of 50 µg/mL, while in panel C, 25 µg/mL was used. Mean oocyst number/mosquito midgut is shown by the red horizontal bar.

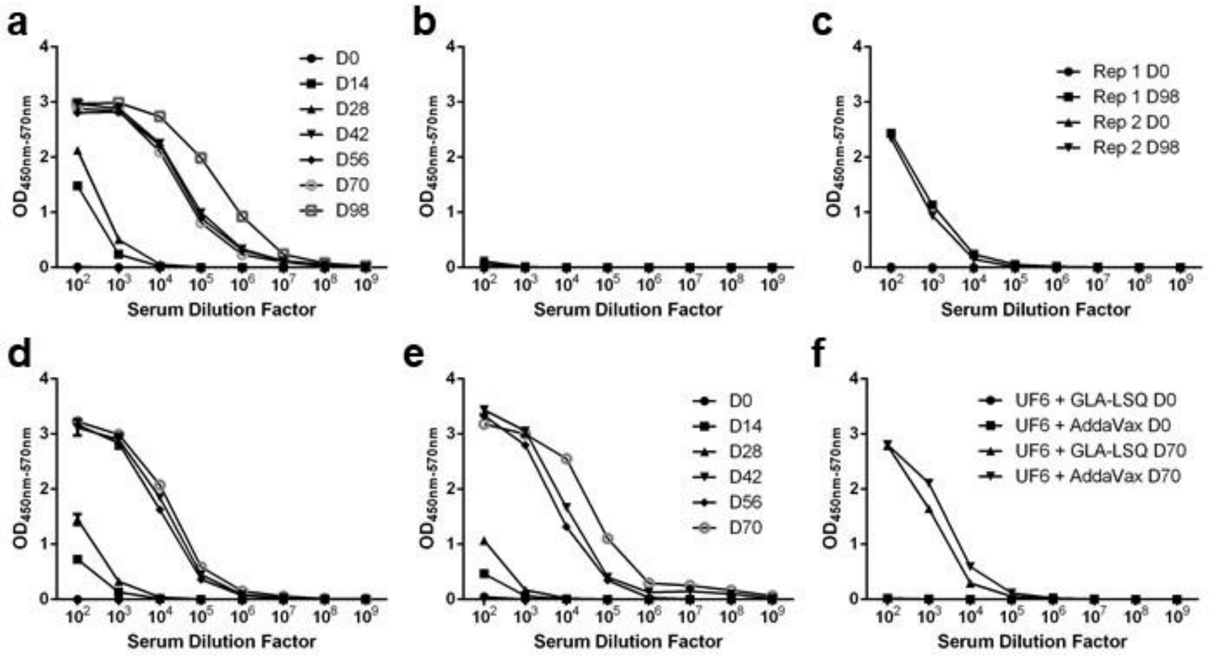


**Fig. S5. Indirect enzyme-linked immunosorbent assay results of replicate mouse studies. (a-c)** A second cohort of mice were immunized (i.m.) with UF6b:GLA-LSQ in a prime and two boost regimen (boosts on D28 and D70) for the 98-Day study (20 µg/dose/mouse). Antibody response to UF6b **(a),** peptide 7 **(b),** and peptide 9 **(c)** were measured. The peptide 9-specific response in the second cohort is plotted along with that from the first cohort of mice tested (Indicated as Rep 2 and Rep 1, respectively). **(d-f)** The 70-Day study was also repeated in which mice were immunized in a prime and two boost regimen (boosts on D28 and D56) (20 µg/dose/mouse) with either UF6b:GLA-LSQ **(d)** or UF6b: AddaVax™ **(e).** The antibody responses to UF6b **(d-e)** and peptide 9 **(f)** were measured. ELISAs were performed on pooled sera collected at designated timepoints. Error bars indicate SEM of triplicates.

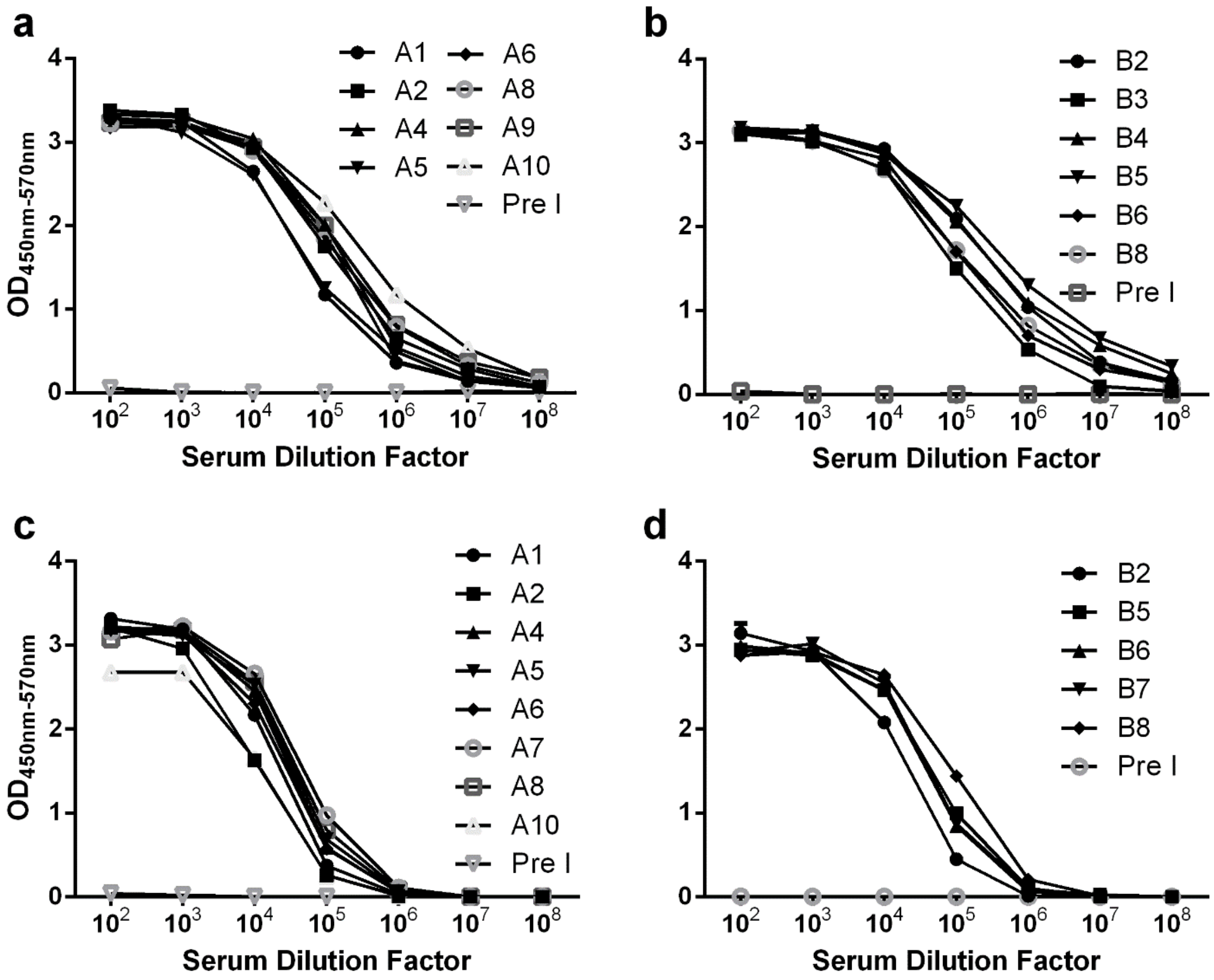


**Fig. S6. Indirect enzyme-linked immunosorbent assay results for individual mice.**

**(a-d)** Mice were immunized in a prime and two boost regimen (boosts on D28 and D56) (20 µg/dose/mouse) with either UF6b:GLA-LSQ **(A,C)** or UF6b:AddaVax™ (**b,d)** in two separate cohorts of mice **(a,b and c,d)**. A1-10, B1-10: mouse identification numbers. Each data point represents an individual mouse. Error bars indicate SEM of triplicates.
